# An agent based model (ABM) to reproduce the boolean logic behaviour of neuronal self organized communities through pulse delay modulation and generation of logic gates

**DOI:** 10.1101/2023.11.10.566538

**Authors:** Luis Irastorza-Valera, José María Benitez, Francisco J. Montáns, Luis Saucedo-Mora

## Abstract

The human brain is arguably the most complex “machine” to ever exist. Its detailed functioning is yet to be fully understood, let alone modeled. Neurological processes have logical signal-processing aspects and biophysical aspects, and both affect the brain structure, functioning and adaptation. Mathematical approaches based on both information and graph theory have been extensively used in an attempt to approximate its biological functioning, along with Artificial Intelligence approaches inspired by its logical functioning. In this article, we present an approach to model some aspects of the brain learning and signal processing, mimicking the metastability and backpropagation found in the real brain while also accounting for neuroplasticity. Several simulations are carried out with this model, to demonstrate how dynamic neuroplasticity, neural inhibition and neurons migration can remodel the brain logical connectivity to syncronize signal processing and obtain target latencies. This work demonstrates the importance of dynamic logical and biophysical remodelling in brain plasticity.

## 1. Introduction

Studying the brain’s structure is a difficult task for multiple reasons, tackled from very different perspectives [1]. There is no univocal model or chart of the brain because of the individual variability, which is not necessarily caused by pathologies. This variability makes the overall description of the brain and its standardization on the micro- and nanoscales, challenging. Furthermore, obtaining measurements through microscopy presents its own difficulties: tissue handling, contrast, stain density, dissection, lighting, etc. [2].

MRI (Magnetic Resonance Imaging) has made the task of studying the brain structure easier, with a greater resolution [3] and less risk than radiation-based methodologies such as X-rays, Computerized Tomography (CT-scans) or Positron Emission Tomography (PET), However MRI is, in general, contraindicated for patients with implants or pacemakers, limiting its applications. The more recent MRI’s functional variant (fMRI) leverages changes in blood flow associated with brain activity, obtaining some promising global brain mappings [4, 5, 6, 7, 8]. It is used prior to surgery and is key in bringing together different time and spatial scales in the brain [9] and so explore feedback and feed-forward behaviour within cortical layer hierarchy [10]. However, the technique is also conditioned by task execution, signal-to-noise ratio and patient comorbidity [11]. There are other techniques to understand the structure and functioning of the brain such as Diffusion Tensor Imaging (DTI) and Transcranial Magnetic Stimulation (TMS).

Zooming into the nano-scale, the brain’s most basic individual parts are neurons: electrically excitable cells allowing for the transmission of information through the nervous system. Neurons have various shapes and have specific functions, but they do usually share a basic structure composed of dendrites conveying information from preceding neurons into a nucleus (soma), which sends signals along an axon – enveloped to varying degrees by a myelin sheath – to the next one. Measuring a neuron activity or inactivity is a complex task which is subject to morphological and method-driven variations. Neuron counting has evolved from microscopical measures [12] – with help of MRI [13], stereology [14, 15] or cytometry [16] – to state-of-the-art solutions involving Deep Learning [17]; whereas tracking individual neuron activity necessarily implies local measurement of biological and/or electrical indicators. Examples of targeted biological indicators are proteins such as PSD-95, which decay is linked to Alzheimer’s disease [18, 19, 20, 21]. Electrical inticators are measured through). Electrical indicators are obtained through electrophysiological studies [22, 23].

This article presents a different approach to study brain functioning and, in particular, damage-related changes in its functioning. The purpose is to replicate the transmission of signals within neurons within the same brain region (or cross-regions, boundary conditions abiding) by programming, considering some simplifications due to the complexity of the real brain. However, with the modeling approach three major brain properties are addresed: metastability, backpropagation and neuroplasticity. Neuron’s nucleus (soma) will be referred as “neuron” from here onward.

## 2. Modelling aspects

In this section we address some aspects of the model to be introduced below.

### 2.1. Proposed neuronal model and communitarian interactions

Neurons are modelled as cells in an agent-based model. This implies that neurons, as agent-based cells, have communitarian behaviour and interactions, and as biological cells have consumption of resources and may migrate. Neurons have also connections between them (synapses) and propagate pulses through other neurons. Signal transmition is modelled assuming the McCulloch and Pitts [24] neuron model, but incorporating modifications to incorporate also the biological point of view. With this approach, the signaling processing can be measured and studied incorporating mathematical tools and concepts like convergence or accuracy, but a biological interpretation of the neuron and the incorporation of biological aspects in the full network, is also possible.

#### 2.1.1. The McCulloch-Pitts neuron model

The McCulloch-Pitts model is the first mathematical model of the signal processing in a biological neuron [24], capable to fire and behave in a similar way than a real neuron. The neuron has connections with a variable number of presynaptic neurons, each one with a different random potential. Once some of the presynaptic neurons have fired, if the average potential received from those neurons is higher than a certain threshold, the neuron will fire; otherwise it will stay latent. The main drawback of the McCulloch-Pitts model is the restriction to produce only binary outputs (fire or not-fire). Thus, typical artificial neural networks enhances the capabilities of the McCulloch-Pitts model through the consideration of variable weights and more ellaborate firing rules.

In this work we implement an enhanced McCulloh-Pitts neuron signal processing model within an agent cell in an agent-based model. This way, neurons process signals, but also are capable of performing plastic remodelling, migrate or inhibit, as its biological counterparts, dynamically changing the brain processing map by biological interactions, not just by signal weight changes.

#### 2.1.2. Inhibition and excitation

In the biological model, there are neurons in charge of supervising the community learning. The neurons in charge of the supervised learning of the community can emit inhibitory and excitatory signals, which are propagated through the network with back-propagation. At each iteration of the model, 0.05% of the neurons with a higher signalling threshold for inhibition or excitation are the ones that in subsequent steps will change their behaviour. In the proposed approach, when a neuron enters inhibition it is modeled by assuming a higher firing threshold, which acts as a switch off interruptor. The excitation mode is modelled through plastic remodelling and by setting a lower firing threshold.

#### 2.1.3. The plastic remodelling process

In the proposed model the plastic remodelling process is addressed through the changes in the connectivity of a given neuron with its pre-synaptic neurons: when a neuron receives enough excitatory signalling, the neuron modifies its pre-synaptic neurons. This is mandated when the neuron in charge of supervising the training process demands to receive a signal earlier than its current latency. Then, the neuron under plastic remodelling looks for new presynaptic neurons between the ones that have fired earlier than its current presynaptic connections.

#### 2.1.4. Migration

In the present model, a third way by which the neurons alter the pulse propagation is migration. New neurons may be added to the model and are initially placed randomly within the brain, making connections with the surrounding neurons. Future model development will include different cell migration criteria to improve the adaptability of the neurons.

### 2.2. Metastability

Different wave bands in the brain (alpha, beta, mu, etc.) may be positively or negatively correlated among themselves [25], but they are nevertheless out of phase and so have to be processed. Metastability explains how the brain coordinates multiple input signals from multiple receptors in time (neural oscillations, i.e. generated by sensory neurons after receiving stimuli) into a coherent, unison response, locked in frequency (like the movement order sent to a muscle from a motor neuron). This way, the brain makes sense of all the unsorted, unorganized information it receives to produce meaningful preprocessed data that will decide a given outcome [26]. These brain waves synchronise, travelling in sequence, and experience nonlinear instabilities [27]. Metastability coordinates the flow of information between brain areas in long spatial and temporal intervals, generating perception, emotion, and ultimately, cognition itself, restarting the latter after sleep [28].

### 2.3. Backpropagation

Neural backpropagation happens from the axon hillock to the dendrites aiming for the dendritic voltage-gated calcium [29] or sodium channels [30] when the soma undergoes depolarization. There is empirical evidence of pyramidal neurons (with separated apical and basal dendrites) performing back-propagation [31] and modelling attempts of this phenomenon are abundant [32, 33, 34]. Nonetheless, this neurological concept differs greatly from its more widely known mathematical counterpart in Machine Learning, as it will be explained in the next subsection.

### 2.4. Plastic remodelling

Neuroplasticity is the brain’s ability to modify its connections and/or paths so the information can get across despite severed or malfunctioning axons, avoiding them and finding alternatives. It has been empirically proven, and it is well known, that this process takes place in brains of any age, regardless of illness [35]. This neuroplastic process can be structural, by regeneration or collateral sprouting (reactive synaptogenesis, rerouting, retraction [36]); and can also be functional, by task relocation within an already existing neural infrastructure (homologous area adaptation, cross-modal reassignment, map expansion, and compensatory masquerade [37]). Structural changes like neurogenesis occur almost exclusively in the hippocampus and olfactory bulb [38] and can be enhanced positively (exercise and good environmental conditions), or negatively (stress, injury, disease). Some functional changes such as map expansion are ubiquitous and life-long [37].

Neuroplasticity is ultimately responsible for the adaptation of the brain’s structural and/or functional connections to external stimuli by senses, and intrinsic stimuli by learning. This adaptation occurs during growth, but also after injuries [39], regaining lost functionalities [40] even to astonishing extents in some cases [41]. Diaschisis (or “functional splitting” [42]) is a phenomenon closely related to neuroplasticity, consisting of a sudden function change/inhibition in a brain area caused by disturbance/damage in another distant but structurally connected zone. This happens mainly in severed connections in the Central Nervous System (CNS).

### 2.5. Biological and Artificial Neural Networks

Although artificial neural networks (ANNs) are in fact inspired by the signal processing of their biological counterparts in the brain [24], several authors [43, 44, 45] have pointed out major differences between them. Thus, the approach presented in this paper is hybrid, introducing Graph Theory and Machine Learning considerations while accounting for real brain phenomena for biomedical accuracy.

In ANNs, the signal propagates forward in the first place. By repeatedly applying the chain rule of derivatives, one can define the rate of change (gradient) of the output prediction *ŷ* or any layer immediate outputs *z_i_* in relation to a given input *x_i_*, being *z_i_* = *W_i_x_i_* + *b_i_*. After evaluation of a given loss function *E* = *f* (*y, ŷ*), backpropagation in ANNs is the process by which signals travel backwards (from outputs to inputs) in order to correct or rearrange previous steps and so approach a target output *y*, correcting the weights *W_i_* and biases *b_i_* of each neuron layer to “learn” the correct combination that yields the target outcome. Using Automatic Differentation (AD) [46], that gradient derivation is carried out by most Artificial Intelligence packages such as Pytorch© or Tensorflow©.

A monotonic (ever-growing) activation function is enforced to cast the layer’s output *a_i_* = *f* (*z_i_*) into a [0,1] interval, such as hyperbolic tangent (tanh), rectified linear units (ReLU), exponential (sigmoid) or any of their variants. In the most common ANN architectures, such as MultiLayer Perceptrons (MLP), Convolutional (CNN) or Recurrent Neural Networks (RNN), layers are usually fully connected, meaning all neurons in layer *i* are connected to all their inputs in the previous layer *i −* 1 and all their outputs in the next one, *i* + 1, and the relevance of connections are typically left to the weights, even though in some deep ANN “weak” connections may be eliminated. There are some exceptions like Graph Neural Networks (GNN), in which partially-connected graphs can still have their nodal and edge attributes updated through specific functions instead of directly computing the gradient, a technique known as “Message Passing” [47].

As for the activation functions in real neurons, membrane activation (responsible for neurons and muscle cells) does have a threshold (an electrical activation potential around −55 mV, which can be graded), but remarkably, its activation curve is not monotonic, going through separate depolarization (*Na*^+^ ions enter, potential rising to the maximum, +40 mV), repolarization (*K*^+^ ions exit, potential decreasing to the minimum), refractory (hyperpolarization) and resting states (potential stabilized at −70 mV). Synapses themselves are regulated by very convoluted biochemical (neurotransmitters, proteins) and electrical processes which are usually not considered in an ANN model at all. On top of that, it takes milliseconds for a single synapse to complete it [43], a somewhat slow rate if compared to some high-performance of multi-layered ANNs after training, which does require longer times, especially in deep networks with convolutions [48].

Conversely, inhibitive and excitative paths for back-propagation in the real brain are distinct, since neurons produce either inhibitory or excitatory synapses, but not both. Real inhibition/excitation paths are diffuse since biological neurons are not fully connected, let alone layer-structured and they receive no information other than their preceding neighbours’ outputs [43], which constitutes a direct obstacle to perform back-propagation where all weights are needed, sometimes referred as the “synaptic assignment problem” [49]. Besides, neurons in primates tend to activate based on attention mechanisms rather than on error back-propagation when reacting to visual stimuli [50]; some ANN models consider this [51]. For example, they active when learning to recognize and classify physical shapes [52].

Synapses between neurons are also subject to plasticity: their intensity and distribution change according to the task [53], electrical stimulation [54], age [55, 56] and damage [36, 57]. In essence, synapses are also trainable [58]. Indeed, data scientists have suggested more realistic computational approaches which accounts for neuroplasticity as a simple given rule [59, 60, 61, 49, 44, 31, 62]. In biological neural networks, the Hebbian learning rule applies: “neurons that fire together, wire together”, implying that functional connectivity determines the structural connectivity, so structural plasticity would submit to functional needs. That poses a problem for a realistic implementation of ANNs, since neural connections have to be strong enough to ease memory recalls but not too strong to create numerical overcharge. Some solutions to this problem imply transient nonlinear analysis [63].

## 3. Methodology

All needed data is produced by self-made code in Python© considering some limitations such as size. Size limitations in the model are needed with because the real amount of neurons is estimated to be around 100 billion [64]. Moreover, the order of magnitude of total synapses in a young brain is 10^15^, even though they change in number and spatial distribution with age [65, 66, 67] and illness [68, 69, 70] and lack thereof [71], specific area and chemical procedures. Indeed, mapping the brain is a difficult topic, as previously explained. In this work, synapses are considered a quasi-instantaneous, purely electrical process. In the proposed model we have used 317,321 synaptic connections and 42 thousand neurons to represent the parts of a logic gate. For comparison, a modern CPU contains roughly 100 millions of logic gates. The proportion between the neurons for the reproduction of the logic gate and the ones needed in a future application to reproduce computing capabilities are of the same order.

Of the neurons used in the model, one thousand are the stimulated neurons, in charge of the initiation of the spikes. Another one thousand neurons are in charge of supervising the learning of the network. The remaining 40 thousand neurons are intermediate neurons that constitute a dynamic network that undergoes remodelling of connections. Mathematically all the neurons are equal, but they have assigned different roles in the network.

A physical domain in R^3^ is created, consisting of a 4*×*0.5*×*0.5 mm prism. The volume is 1 mm^3^ which reproduces the estimated neuronal density of the brain of 40, 000 neurons*/* mm^3^ [72]. In two boundaries of the domain (frontal and posterior faces of the prism), *n*_1_ stimulated and *n*_2_ supervising neurons are placed. Between these two areas, the number is much higher, *n*_0_ = 20 *×* (*n*_1_ + *n*_2_) neurons. The spatial coordinates of these neurons are given randomly within the domain, in an sparse pattern. All these neurons represent different global inputs (emitter neurons) and outputs (receptor neurons) their synapses constitute the processing paths between them in a certain regionof the brain.

The initial random synaptic connections are deployed between the neighbouring neurons. Neighbouring connections (neuron spatial map) is obtained through Delaunay’s triangulation, generalized to N-dimensions through the Bowyer-Watson’s algorithm [73, 74]. This approach is widely used to create unstructured meshes [75], and creates tetrahedra whose vertices are contained within their correspondent circumsphere. Its homogenizing properties in joint angle and edge (axon) length are useful to accurately represent a network composed of one specific type of neuron, morphologically similar though not exactly identical. This configuration also avoids unforeseen elements in the structure (unwanted neurons or connections between them), since it prevents edge intersection. After this process, the initial structural connectome (the brain’s infrastructure) is established.

Every neuron *i* is given an initial weight *w_i_ ∈* [*−*1, 1] and an activation threshold *α_i_* (initially zero), so that if *w̅*ℐ*_i_* ≥ *α_i_*, it will be activated, being ℐ*_i_* the set of input neurons of *i* fired at this time step, and *w̅*ℐ*_i_* its average value. An initial (structural) connectivity is assigned as well. In a first approach, structural and functional connectivity were considered equivalent, so information would flow always from emitters to receptors. Nonetheless, neurons are not aligned in the real brain, neither sense-wise (soma to telodendria) nor path-wise (beginning to end), so the sense is randomly decided for each of them. The only rule that applies is the non-connectivity of input or output neurons among themselves if they are treated as stimulated of supervising neurons respectively. Following this reasoning, whichever inputs a given soma (nucleus) receives are considered its dendrites, and so its outputs would correspond to its axon terminals (telodendria), as in multipolar neurons all around the Central Nervous System. Then, edges in this model represent either dendrites or axon terminals, whereas nodes include every other part in the middle (nuclei, axon hillock, axon, myelin sheath, etc.).

For emulating brain metastability, the density distribution of active supervising (or output) neurons must synchronise with a given desired target signal, for instance, an exponential curve centred at a certain propagation step. To train that synchronization, certain sets (fractions) of input neurons (stimulated, *n*_1_) are sequentially lit by default during the initial time steps and so the firing of neurons propagates forward eventually reaching the supervising neurons and activating some of them. At each step, inhibitory and excitatory signals propagate backwards to neurons with incorrect status, inhibiting the neurons that should had been off but are on and exciting the inactive ones that should have been active. This is the supervised learning mechanism that the *n*_2_ neurons apply to the neuronal net. After this first training iteration, newly-corrected forward propagations are sent to increase accuracy with each step. By the end of a simulation, the more closely the synchronised combined signals approach the target distribution, the more accurate the model will be.

Due to the sense randomness introduced, some signals are effectively lost, as expected, reaching a point where they cannot find a viable path matching the activation conditions and so they disappear. This can be interpreted as a normal consequence of network randomness. Another interpretation is that of signals travelling to other out-of-scope regions of the brain (far-away connectivities). They also account for neuroplasticity, allowing for new paths to be explored in different simulations. To further enhance signal synchronisation, a specific counter is set so that if the hop-distance (number of connections traversed) between two neurons is equal to the difference in time steps, the activation of the precedent one is enhanced.

A specifically built function introduces a percentage of neurons changing status. The purpose is to mimick functional neuroplasticity by searching for the most busy neurons (through which most signal paths go through) and exciting or inhibiting a small set of them accordingly. This excitation/inhibition is modelled by decreasing or increasing their activation potentials and through plastic remodelling. This remodeling depends on whether those paths are mostly inhibitive or excitatory, and recreates the variance in neurotransmitter receptors happening in synaptic plasticity [76]. In Graph Theory terms, these neurons (or nodes) are central, since they have the highest degrees (number of neighbours). The fraction of neurons affected by neural plasticity must be limited to *<* 0,05 % of the total number (i.e. 20) at each adaptation step to avoid numerical instabilities, as previously explained.

As for structural neuroplasticity, that is, changing the connectome’s infrastructure by rearranging, severing and/or creating axons; certain damaged neurons over time (due to age, illness, injury or any combination of them) could force reactive synaptogenesis to occur, relocating connections either in the damaged neuron’s neighbourhood (easiest, most straight-forward) or in unexpected far-away places (emulating axon rerouting and/or age-induced density loss). One way of doing that comes in response to backpropagation: negative-weighted neurons could reconnect along inhibitory paths and positive-weighted neurons could do likewise for excitatory ones, reinforcing them. That way, each possibly useful pathway is optimize while the rest perish if unneeded. If neural damage does not take place, axon retraction could be put into practice by progressively trimming (a fraction of) the least used connections for global connectome efficiency - a process commonly known as “pruning” in Neuroscience [77].

Also, the neuronal migration is implemented in the model. In this case, new neurons are randomly appearing in the net and connecting with the transition neurons. This appearance is also limited to 0.05% of the total number of neurons at each adaptation step.

The signal initiates with a stimulation of the *n*_1_ neurons in different steps. At the beginning of the calculation 25% of the *n*_1_ neurons are fired, in the next propagation step other 50% of those neurons are fired, and in following one, the other 25%. So when the last *n*_1_ neurons are fired, the signal originated with the first stimulated ones are 2 propagation steps ahead. Then, the signal is calculated through all the propagation steps from the fire of the *n*_1_ neurons to the fire of the *n*_2_. The network is self-remodelled, and a new cycle starts with a new propagation.

### 3.1. Creation of logic gates with neurons: modification of the McCulloch-Pitts boolean model

The model shown in the previous section is used in the squares of the Figure 1, as *N*_1_ and *N*_2_. Their role is that given two stimuli *I*_1_ and *I*_2_, not necessarily coordinated, those will be synchronized through the networks *N*_1_ and *N*_2_ as validated before. This section is intended to study boolean configurations where this model can reproduce logic gates from asynchronous stimuli. The boolean analogy is fire (true) and not fire (false). Figure 1 shows 3 schemes for AND, OR and NOT gates.

**Figure 1:**
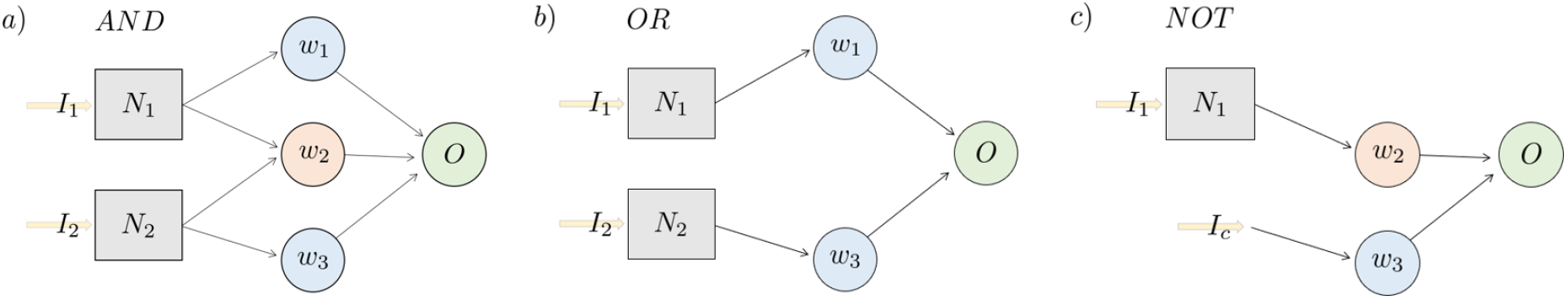
Schemes of logic gates for asynchronous stimuli using the methodology proposed.

Those gates, with their elements, are explained below.

#### AND gate

In this case *w*_1_ *>* 0, *w*_3_ = *w*_1_, *w*_2_ *< −w*_1_, and the *α* of the neuron *O* is 0. Si, if *I*_1_ or *I*_2_ are stimulated alone, the average *w̅* is lower than the threshold *α*, only if both are fired at the same time the *w̅ > α*. Of course, any other combination of values for those variables can achieve the same objective as far as those proportions are fulfilled.

The boolean output (*O*) can be defined as *O* = *I*_1_ *AND I*_2_, if there is a stimulation in *I*_1_ and *I*_2_ the output in *O* is to fire, otherwise *O* would not fire.

#### OR gate

In this gate any stimulus in *I*_1_ or *I*_2_ will fire *O*, since the inhibitory intermediate neuron with *w*_2_ is removed. The boolean output (*O*) can be defined as *O* = *I*_1_ *or I*_2_ in this case.

#### NOT gate

This logic gate is intended to switch between the state of *I*_1_ and *O*. Here only one input is required, but a continuous fire *I_c_*is needed to ensure that when *I*_1_ is not stimulated *O* is. So, for this reason a constant firing signal is needed. And when *I*_1_ is stimulated, the inhibitory neuron *w*_2_ is activated and so *O* is not fired, since now *w̅ < α*. The boolean output (*O*) can be defined as *O* = *NOT I*_1_.

## 4. Results

This section shows the results of the implemented model. Regarding computational times, in a serial code in Python, run in a single core, it took 43 seconds. For every adaptation step of the neuronal network, 24 iterations were needed in total to adapt the network in the two cases. As mentioned before, the adaptation process was done through two processes, a logical and a biophysical one. The logical one consists of inhibition and excitations that results in plastic remodelling. The biophysical one consists of migration of new neurons to the area. The results show the capability of the network to adapt to change the delay of a signal, which is essential for the logic gate models proposed.

### 4.1. Decrease and increase the latency of a pulse

In this case, with the model described in the methodology, the delay of a pulse is increased and decreased from the reference latency of the network. The reference latency is 80 propagation steps (case 1 of Figure 2, and the target is to reduce it to 40 propagation steps (case 2) and to increase it to 120 propagation steps (case 3). Considering that the length of the prism studied is 4 mm, and that it has a reference latency of 80 propagation steps, the signal advances as an average 0.05 mm for each propagation step. The speed of the signal in the brain can range from 0.5 to 100 m*/*s, with implies that considering, for example, 10 m*/*s as the reference velocity, each propagation step will be equal to 5 *µ*s.

**Figure 2:**
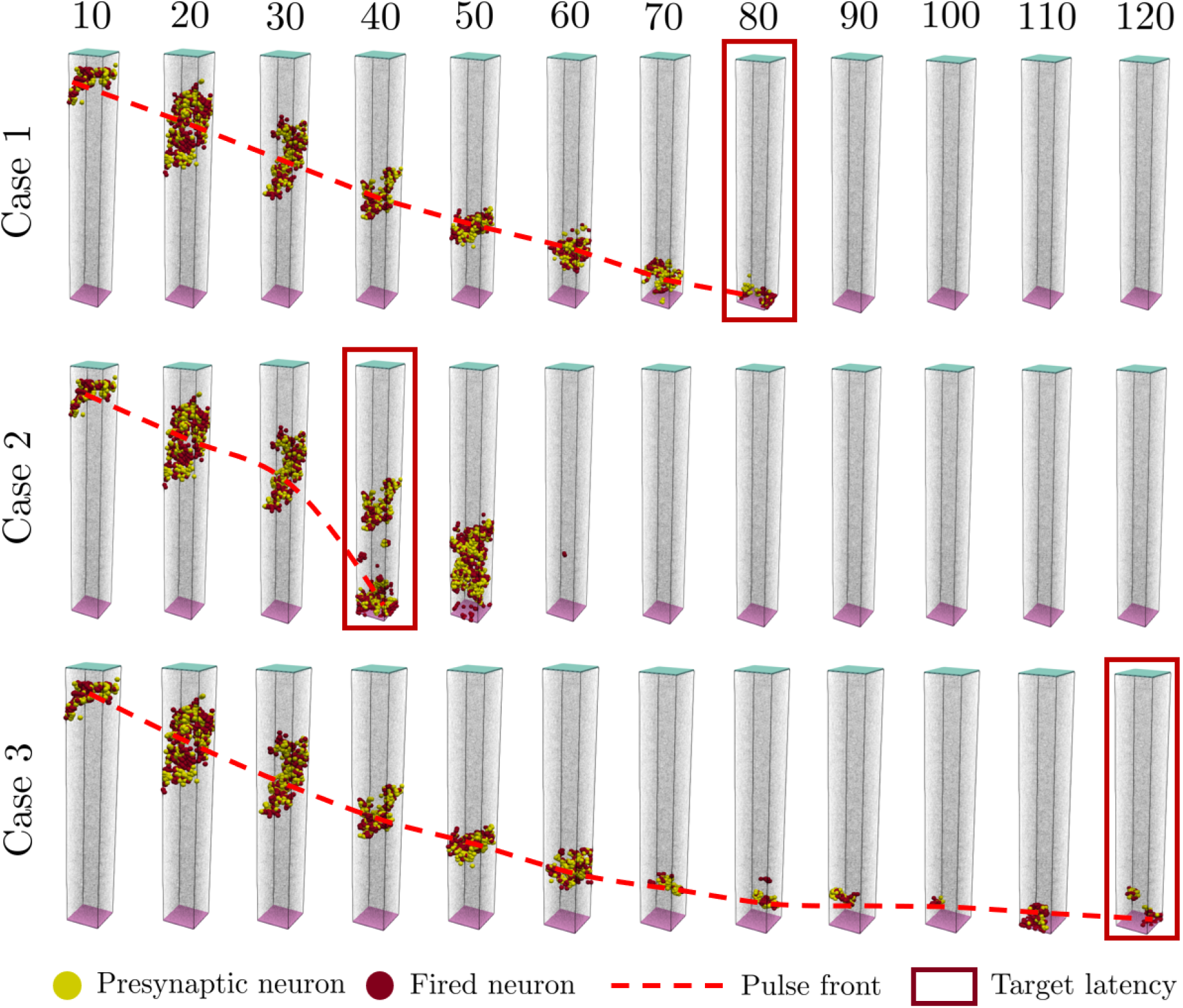
Signal progression in the 3 cases studied: reference (Case 1: signal arrival takes place at 80 propagation steps), reduced delay (Case 2: signal arrival takes place at 40 propagation steps) and increased delay (Case 3: signal arrival takes place at 120 propagation steps). Note that the reference case has an almost constant propagation rate, whereas the other two cases modify the signal propagation rates.

In Figure 2 the top square shown in every lapse (coloured in light blue) are the neurons excited with the predefined input signal. The bottom square (coloured in light purple), is where the supervising neurons are. So the signal goes from the top to the bottom, and once the supervising neurons get excited, they send the backpropagation signals according to their target signal delay.

Figure 2 shows the firing steps of the adapted networks. The case 1 shows the firing steps of the reference network, the one generated randomly as explained in the methodology. This case 1 is also the starting point for the remodelling of the network to achieve the states of Case 2 and Case 3. Case 2 is the result after the remodelling to reduce the latency of the network. Also in Figure 2, the red dashed line shows that the front progress velocity is not constant, at the end of Case 2 it is accelerated, and at the end of Case 3 the velocity is reduced. In the Case 1 this propagation velocity is roughly constant as a consequence of the equally random distribution of the neurons and their synapses.

In Case 2 of Figure 2, at the propagation step 50 it can be seen that the signal front goes backwards and intercepts other signal front in the neuronal network. Those are spurious signals that are naturally diminished in the model before reaching the output.

The results shown in Figure 2 have been obtained after the remodelling process. Figure 3, shows the changes involved in the inhibition and excitation needed to change the delay of the signal for Cases 2 and 3. In the Case 2 of Figure 3 the plastic remodelling of the neurons allow the signal to go faster through the network. As mentioned in the methodology, the plastic remodelling is done by changing the synapses. The red dashed line shows that as the plastic remodelling progress, the synaptic connections are done at a higher distance. The Case 3 of Figure 3 shows the remodelling carried out by the network to increase the latency of the system. Here some neurons are inhibited to increase the path of the signal, and other neurons migrate and connect with the network to prevent the cut of the signal, and ensure its continuity.

**Figure 3:**
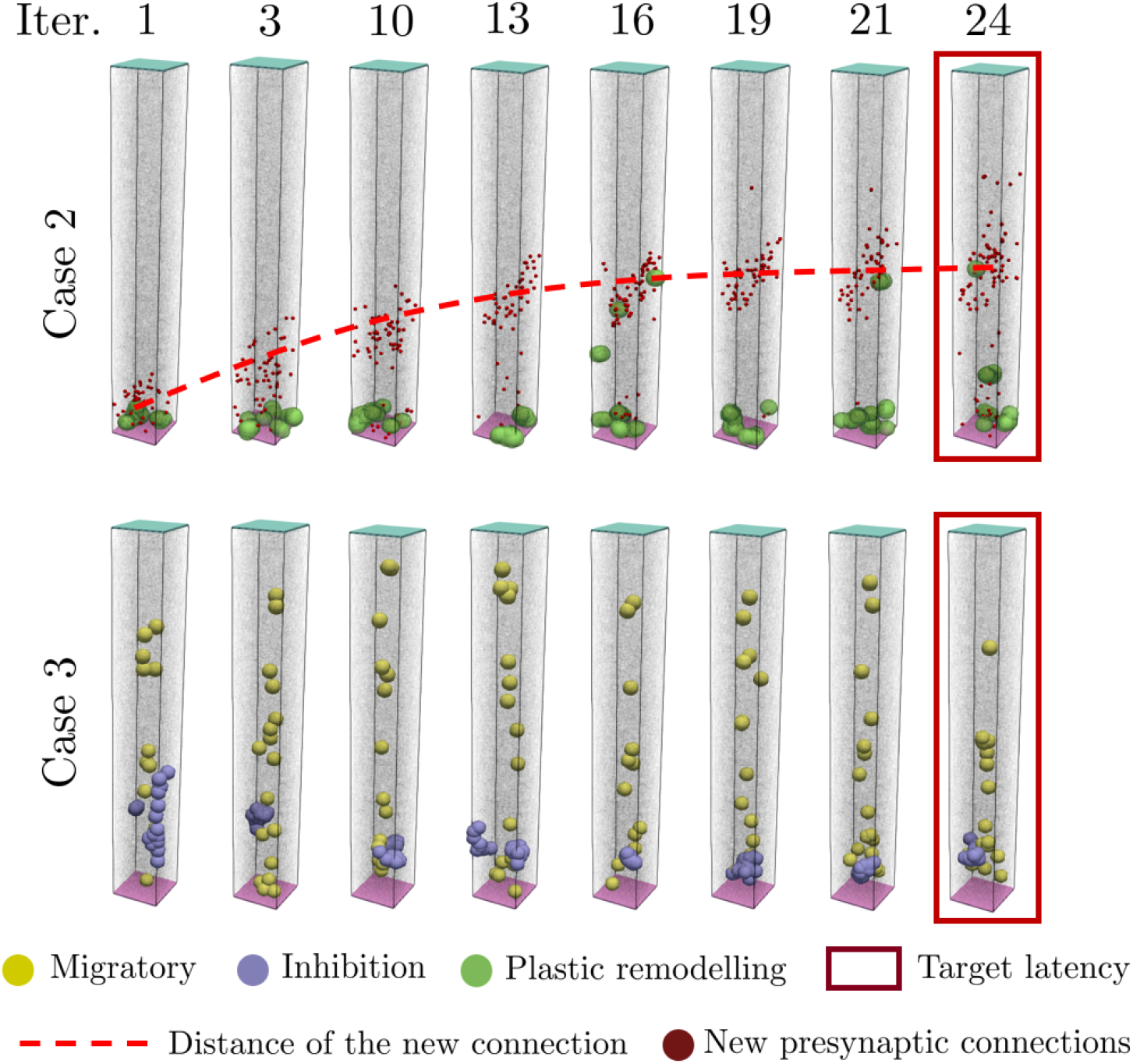
Remodelling strategies adopted by the neurons for cases 2 and 3 during progressive iterations to achieve the target latency. In Case 2 it is observed that changes are mainly due to plastic remodelling (changes in synaptic connections, new presynaptic connections), and with every iteration the new and the average distance of the connected neurons increase; after a given number of iterations the neural structure converges. In Case 3 synaptic plastic remodelling seems to be not sufficient or adequate, so neurons enter in inhibition or migratory status to achieve a delay in the requested latency of the signal mandated by supervisory neurons.

In quantitative terms, Figure 4a shows the evolution of the delay of the signal for different remodelling iterations until they reach the target delay. Figure 4b shows the output signal at different remodelling iterations for both the increased and decreased delay.

**Figure 4:**
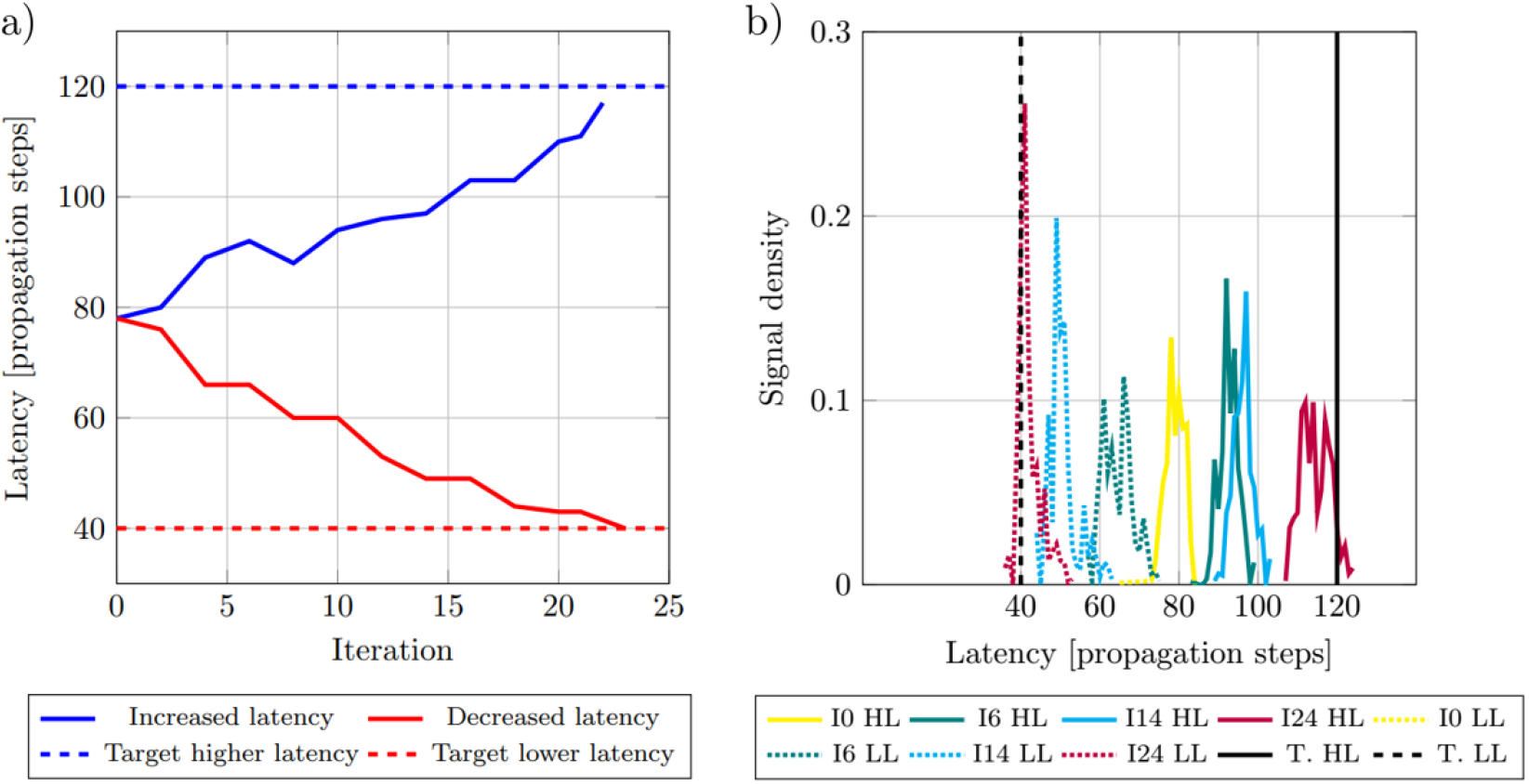
Evolution of the output signal with the remodelling iterations. Left: evolution of the latency of the signal towards the target with iterations. Right: Evolution of the received signal density with propagation steps, during the iterations. Right legend: HL (continuous curves) stands for higher latency target case, LL (dashed curves) stands for lower latency target case, I0 (yellow) to I24 (red) stand for iteration number and T.HL and T.LL stand for target higher latency and target lower latency, respectively.

This proves that with two of those prisms in parallel, different uncoordinated stimuli originated in different parts can result in a coordinated pair of signals capable to be used for logic calculations. This is shown in the next section.

## 5. Discussion

The results reflect a simplified yet sufficiently realistic representation of the brain’s information flow: it imitates metastability by coordinating one or multiple diverse and out-of-phase input signals into a single output target signal, while enforcing biologically plausible backpropagation with separate inhibitory and excitatory paths - further enhanced by the introduced neuroplasticity functions. It must be mentioned that this model reaches the target by organically adjusting functional and structural paths, without any explicit optimization parameters nor loss function. In spite of these satisfactory results, the authors are aware of this model’s limitations, especially regarding the step around which the target signal is centered.

On the one hand, moving the target distribution to the left (fewer steps) is equivalent to organically shortening paths to arrive to the same output. Computationally speaking and for a negatively-weighted, undirected partially connected graph like this, this problem can be solved by algorithms such as Bellman-Ford’s [78], Johnson’s [79] or Floyd-Warshall’s ([80], worst-case complexity O(*V* ^3^), being *V* = *n*_0_ + *n*_1_ + *n*_2_ the total number of connected vertices). Obviously, it is unlikely that this network heuristically solves such a problem in a limited amount of steps.

On the other hand, should the target move “rightwards” (more steps), the model would have to artificially extend paths to synchronise the signals, searching for the longest path, which is a NP-hard problem, even NP-complete if a certain length is sought. Some solutions exist for directed, acyclic graphs [81, 82] (even with perturbations - adding and eliminating edges - [83]), but checking whether a Delaunay-generated graph contains cycles is a NP-complete problem itself. [84, 85].

To address this issue, some global (clustering, small-world coefficients) and node-dependent (vulnerability, shortest path) graph parameters could be introduced and lever-aged to choose the most convenient path - short or long, depending on the target signal’s position in time. They would deliver useful information on the topological characteristics of the connectome, so that certain clustered areas –known as “hubs” [86]– could be avoided (shorter paths) or crossed (longer paths), whereas vulnerability-related indicators could provide very valuable information on which alternative paths to follow when certain connections (structural or functional) are damage or even severed - neuroplasticity in practice. Such parameters have already been studied on a brain regional level [87, 88, 89, 90], but not on a neuron level like suggested here. A graph-based approach entails two self-evident ramifications for brain networks: the multi-scale approach - in both time [91] and space [92] and the involvement of Graph Neural Networks [93, 94, 95, 96, 97] or similarly-flavoured neural network techniques [98, 99] - notwithstanding the aforementioned caveats.

Another option would be to organically shorten or enlarge neural pathways if the target gets closer or further, respectfully, although biological evidence for such a behaviour remains elusive. Introducing time constraints, such as refractory periods [59] replicating membrane functioning could help in this goal, since the present approach does not consider real time but propagation steps. In this scenario, neuron length does not play a practical role, so the only meaningful way to shorten (or enlarging) neural paths would be to artificially skip (or add) synapses along the way.

The next significant step to be taken is the development of a bio-mechanical model of the brain which reflects the interaction between information transmission and external loads or accelerations (leading to Traumatic Brain Injury), loss or damage of axons (neurodegenerative diseases) or even areas with distinct properties (tumor growth, brain stiffening caused by Alzheimer’s). There is a plethora of bibliography proposing bio-mechanical models [100, 101, 102, 103, 104, 105, 106, 107, 108, 109], but the correlation between physical damage and information flow has yet to be analytically described. Also, the model itself could be enriched by including and emphasizing the role of other parts of the neuron (like the myelin sheath) or another components of the CNS, such as glial cells [110].

Bearing all these suggestions in mind, futher developments of this code are being addressed, aiming at an even better-performing model with bio-mechanical implications.

## 6. Conclusions

In this work some aspects of the modeling of the plasticity of the brain have been addressed. Remodeling of the brain have both logical (presynaptic connections and signal processing strengths) and biophysical aspects (cell migration, community behavior, inhibition and excitation, among others). The relevance of these aspects have been addressed in brain plasticity, in particular in brain remodeling to reach target signal latencies. The results are promising and motivate further research to better understand changes in brain processing, like in aging, illness and injuries.

The present model is also capable to reproduce the boolean logic behaviour of neuronal communities. This is done with the biological rules for interaction, and considering the neurons as cells capable to perform synaptic connections to transfer information through signalling. This capability of the model is validated with the two examples where the latency of a signal through a neuronal community is delayed or increased.

## Acknowledgements

The authors would like to thank Professor Michel Destrade (Chair of Applied Mathematics, University of Galway) and Sairam Pamulaparthi Venkata (Early-Stage Researcher 8, XS-Meta Project, University of Galway) for their insights on brain modeling as a soft, visco-elastic material, as well as Itziar Terradillos Irastorza (PhD in Neuroscience 2021, UPV/EHU), Hanoi Iván Guillermo Montiel (PhD student, Institut Pasteur) and Edgar Soria-Gόmez (Neuroscience Department at UPV/EHU) for their medical guidance.

